# Functional interpretation, cataloging, and analysis of 1,341 known and new glucose-6-phosphate dehydrogenase variants

**DOI:** 10.1101/2022.09.14.508023

**Authors:** Renee C. Geck, Nicholas R. Powell, Maitreya J. Dunham

## Abstract

Glucose-6-Phosphate Dehydrogenase (G6PD) deficiency affects over 500 million individuals who can experience anemia in response to oxidative stressors such as certain foods and drugs. Recently, the World Health Organization (WHO) called for revisiting G6PD variant classification as a priority to implement genetic medicine in low- and middle-income countries. Towards this goal, we sought to collect reports of G6PD variants and provide interpretations. We identified 1,341 G6PD variants in population and clinical databases. Using the ACMG standards and guidelines for the interpretation of sequence variants, we provided interpretations for 268 variants, including 186 variants that were not reported or of uncertain significance in ClinVar, bringing the total number of variants with non-conflicting interpretations to 400. For 414 variants with functional or clinical data, we analyzed associations between activity, stability, and current classification systems, including the new 2022 WHO Classification. We corroborated known challenges with classification systems, including phenotypic variation, emphasizing the importance of comparing variant effects across patients and studies. Biobank data made available by All of Us illustrate the benefit of large-scale sequencing and phenotyping by adding additional support connecting variants to G6PD-deficient anemia. By leveraging available data and interpretation guidelines, we created a repository for information on G6PD variants and nearly doubled the number of variants with clinical interpretations. These tools enable better interpretation of G6PD variants for the implementation of genetic medicine.

## INTRODUCTION

Glucose-6-Phosphate Dehydrogenase (G6PD) deficiency is the most common enzymopathy worldwide, affecting over 500 million individuals.^1^ G6PD is important in red blood cells since it is the sole source of NADPH needed for detoxification of reactive oxygen species.^2^ Individuals with G6PD deficiency have variants with decreased activity, which can lead to three main clinical manifestations: neonatal jaundice, chronic non-spherocytic hemolytic anemia (CNSHA), and acute hemolytic anemia (AHA) in response to stressors such as certain foods, antibiotics, antimalarials, and infections that elevate reactive oxygen species.^3,4^ Underlying G6PD deficiency is great genetic diversity, with hundreds of identified variant alleles, mostly missense variants in the coding sequence.^1^ Interpreting the function and clinical effects of G6PD variants is critical to prevent adverse drug reactions, which are avoidable by prescribing alternate drugs, and to promote neonatal health by prompting increased monitoring.^3,5^

The G6PD gene is located on the X chromosome, so variant phenotypes are most clearly observed in hemizygotes and homozygotes; heterozygotes are largely unaffected, though phenotypes vary between individuals and over time due to random X-inactivation^4^, but identification is important to avoid anemia and jaundice in hemizygous fetuses.^1,5^ Some G6PD variants reach high frequency in specific populations since they provide protection from severe malarial infection and thus are under balancing selection.^2,6^ Many of these G6PD variants are well-studied and have been reported in dozens of publications along with their activity, clinical manifestations, and distribution across local and global populations.^2^ However, many variants are rare and have little data or conflicting reports on their connection to G6PD deficiency and anemia.

To aid in understanding the effects of G6PD variants, Yoshida and colleagues suggested a five-category classification system in 1971, based on variant activity and severity of anemia.^7^ The classification scheme, called the World Health Organization (WHO) Classification, was modified in 1985 to contain four classes: Class I variants with very low activity resulting in CNSHA, Class II with very low and severe deficiency, Class III with decreased activity and moderate to mild deficiency, and Class IV with near normal activity.^8^ However, even the original authors cautioned that the classification groups were for “convenience” and cutoffs made “somewhat arbitrarily.”^7^ The identification of variants associated with CNSHA that have activity over the 10% cutoff, and the overlap of clinical presentations between Class II and III, led to blurring of distinctions between classes.^9^ Additionally, many variants were classified based on activity or clinical presentation measured in only one individual.^9^

Due to these issues, in 2019 the WHO issued a call to reconsider the G6PD classification system to standardize reporting of activity measurements.^9^ Updated guidelines were released in 2022 to account for variability by requiring activity measurements in at least three unrelated hemizygotes and setting new thresholds for median activity to create new classes: A, variants leading to CNSHA with under 20% activity; B, AHA with less than 45% activity; C, no hemolysis and 60-150% activity.^10^ However, these new guidelines were developed and tested on only 17 variants, and any that do not conform to these classes are grouped as Class U, uncertain.

The WHO Genomics Initiative also identified G6PD classification as a priority for implementation of genetics and genomics medicine in low- and middle-income countries.^9^ The Clinical Pharmacogenetics Implementation Consortium (CPIC) recently released updated guidelines for medication use in G6PD-deficient individuals, including seven drugs with high risk of inducing AHA in G6PD-deficient patients.^4^ To improve genetic assessment of G6PD deficiency and risk of AHA, we sought to collect all reports of G6PD variant activity measurements and clinical presentation. We also tested the implementation of the 2022 WHO classification system, and applied the American College of Medical Genetics (ACMG) clinical guidelines to incorporate data from multiple patients across studies to support clinical interpretation of G6PD variants. Here we provide interpretations for 186 variants previously of uncertain significance or not reported on ClinVar, and a collection of information on variant activity, stability, and clinical presentation. We found that biobank data in All of Us captures diverse G6PD variation in sequence and activity, and aligns with interpretations from previous variant reports. These interpretations can increase the applicability of genetic medicine to individuals with rare G6PD variants.

## MATERIALS AND METHODS

### Variant curation

Variants were curated from the following databases and PubMed searches: CPIC (last updated February 25, 2021)^11^, ClinVar^12^, dbSNP^13^, gnomAD (v2.1.1 and v3.1.1)^14^, LOVD (v3.0 build 26c)^15^, bravo (TOPMed Freeze 8) (bravo.sph.umich.edu), HGMD^16^, and All of Us (see section below). Intronic variants were only included if a clinical annotation (listed on ClinVar) or functional annotation (activity or stability) was provided. Variants synthesized in vitro but not yet identified in patients were listed in Supplemental Table 3 but not Supplemental Table 1 or included in total variant counts. Final database searches were conducted on August 12, 2022.

Variants were denoted by nucleotide change on the negative strand of hg19/GRch37 and cDNA position for transcript NM_ 001042351; cDNA position is reported throughout the manuscript. The most common name for each variant is reported in Supplemental Table 1, but all alternate names are also included in Supplemental Table 2. Variant types were determined by nucleotide changes in the coding region (e.g. single missense, synonymous), and noncoding variant types by VEP annotation listed in gnomAD v2.1.1. Alternate exons from NM_000402 and NM_001360016, and the 5’- and 3’-UTR of NM_001042351, were confirmed using the NCBI Genome Data Viewer.^17^

The only notable corrections made when adding variants to the list were as follows: ClinVar contains two entries for G6PD Puerto Limon on two different transcripts (variation IDs 10381 and 804134), so we listed the variant only once. Variant c.463C>G was presented as G6PD Acrokorinthos though it contained only one of the two nucleotide changes found in the initial characterization of G6PD Acrokorinthos; thus we labeled it as “Acrokorinthos single”.^18,19^ A report of c.1186C>T referenced a case study on the same patients which listed the variant as c.1186C>G, which matched the residue change reported in both studies so thus the data are used as support for c.1186C>G.^20,21^ ClinVar variants 1237186 and 1280478 were not included since they do not encode a nucleotide change.

### Classifying variants according to ACMG guidelines

Criteria provided by the American College of Medical Genetics (ACMG) were applied to reports of curated variants (Supplemental Table 4).^22^ Scores were computed according to a Bayesian classifier with Prior_P 0.1 and translated to interpretations.^23^

Evidence codes PVS1, PS1, PS2, PS3, PS4, PM1, PM2, PM4, PM5, PM6, PP1, PP3, PP4, and PP5 were used in support of pathogenicity. Reports of decreased activity by clinical testing in red blood cells were considered evidence for PS3 (functional study supports damaging effect); reports of decreased activity for variants expressed in model systems were decreased to moderate support (PS3_M). As recommended for interpreting prevalence in affected individuals compared to controls (PS4), reports of variants in unrelated individuals were considered moderate support since case-control studies are rare (PS4_M).^22^ The critical domains used for PM1 (located in critical functional domain) were the NADP binding domain (GxxGDLA) from residues 38-44, substrate binding domain from residues 198-206, dimerization domain from residues 380-425, and structural NADP binding site from residues 488-489.^15,24–26^ Other residues shown to be critical for functional domains in individual variant publications were also included with appropriate references. PM2 (absent or extremely low frequency in controls) was determined using gnomAD v2.1.1 and v3.1.1, with a conservative cutoff of 0.001 for below carrier frequency since the global prevalence estimate is 0.024-0.079 (average 0.05), but allele frequency can be below 0.01 depending on the region.^27,28^ For variants with multiple nucleotide alterations, the lowest frequency site was used to determine if the complex variant frequency was likely below carrier frequency.

Evidence codes BA1, BS2, BS3, BP4, BP5, and BP6 were used in support of benign classification. BA1 (>5% frequency in controls) was determined using gnomAD v2.1.1 and v3.1.1; BS1 (frequency in controls greater than expected) was not used since variants over the average carrier frequency of approximately 0.05 were already captured by BA1.^27^ Reports of normal activity by clinical testing in red blood cells of hemizygotes were considered evidence for BS3 (functional study shows no damaging effect). If the only evidence supporting a pathogenic classification was PM2 (absent or extremely low frequency in controls) but other evidence supported a benign classification, PM2 was disregarded as recommended by previous application of the guidelines.^29^

### Current variant classifications

World Health Organization classifications by 1985 guidelines were taken from published variant lists or individual publications on the variant.^1,15,30^ In rare cases when multiple classes were proposed, the lower was listed. WHO 2022 criteria were applied to classify variants with activity measurements in at least three unrelated individuals by computing weighted median activity and most severe clinical presentation.^10^

ClinVar classifications are given as the interpretation listed for each variant as of August 12, 2022; review status (from 0-4 stars with increasing support, though no G6PD variants had three or four stars) was noted at the same time. “Conflicting interpretations of pathogenicity” was shortened to “conflicting.” If two nonconflicting interpretations were given (e.g. “benign / likely benign”), only the first was listed.

### Variant activity and stability

All reports that measured variant activity are listed in Supplemental Table 3, including activity of variants expressed in model systems. To compare average variant activities, only reports of activity measured from red blood cells of hemizygotes were considered. If not computed by the study authors, average activity compared to the commonly used reference B variant was calculated using the provided normal range in the same study, or in rare cases the normal range from a previous publication from the same laboratory. Average variant activity reported in Supplemental Table 1 was calculated as average across studies weighted by number of hemizygotes with measured activity per study.

All reports that measured variant stability are listed in Supplemental Table 3. A consensus stability for each variant across studies is listed in Supplemental Table 1; in case of conflicts, stability measured in patient red blood cells was reported, or “conflicting” listed if reports from patient cells differed.

### Clinical presentations

The most severe clinical presentation observed for each variant is reported by study in Supplemental Table 1. In decreasing severity, they are CNSHA, AHA, and asymptomatic deficiency. Nondeficiency was also recorded. In cases where patients were reported as deficient without testing for anemia, the presentation was simply listed as “deficient”; this may also include other presentations such as jaundice without anemia, which are reported in Supplemental Table 3.

### All of Us

All of Us data (allofus.nih.gov) were queried using the All of Us researcher workbench cloud computing environment using Python and R coding languages. To find novel variants that have not been previously reported in humans, we used the online All of Us data browser to identify variants in the G6PD gene in the full whole genome sequencing (WGS) cohort. 118 non-intronic variants (considering exons of NM_ 001042351, NM_000402 and NM_001360016) were identified that had not been found in previous PubMed or database searches as detailed above.

A cohort was created for participants with WGS results available and a G6PD deficiency diagnosis with or without anemia (“deficiency of glucose-6-phosphate dehydrogenase” or “glucose-6-phosphate dehydrogenase deficiency anemia”) resulting in 29 participants. Genomic data were extracted from variant call format (VCF) files available to All of Us researchers and loaded into the workspace notebook using Hail and filtered to the G6PD gene region (ChrX:154527010-154552570, GRCh38). Variants comprising multiple nucleotide changes (multiallelic variants) were split and the resulting Hail matrix was written to Plink format. The Plink files were converted to .raw format for upload into R. 101 variants were identified among these 29 people (Supplementary Table 5); for Table 1 the list was then filtered to missense variants. Variant phasing for G6PD was done using Aldy version 4.1.^31,32^ When multiple phasing results were returned, we chose the haplotype that matched the more commonly observed LD structure. For example, for a small number of individual(s) Aldy considered that the variants c.202G>A and c.376A>G were either on the same or separate alleles, however in these instances we chose the haplotype option with these variants on the same allele because that is more commonly observed.

**Table 1:** Missense variants associated with G6PD deficiency in All of Us.

| missense variants | number of participants with deficiency without anemia | number of participants with deficiency with anemia |
| --- | --- | --- |
| none | ≥ 1 | ≥ 1 |
| hemizygous<br>c.[202G>A;376A>G] | ≥ 1 | ≥ 1 |
| homozygous<br>c.[202G>A;376A>G] | 0 | ≥ 1 |
| heterozygous<br>c.[202G>A;376A>G] | ≥ 1 | ≥ 1 |
| c.[202G>A;376A>G]; [376A>G] | 0 | ≥ 1 |
| heterozygous c.376A>G | ≥ 1 | 0 |
| hemizygous c.563C>T | 0 | ≥ 1 |
| heterozygous c.563C>T | ≥ 1 | 0 |
| c.[376A>G;968T>C]; [376A>G] | 0 | ≥ 1 |

Missense variants found in 29 individuals with the code “Deficiency of glucose-6-phosphate dehydrogenase” or “Glucose-6-phosphate dehydrogenase deficiency anemia” in All of Us. All variants found in fewer than 20 individuals, therefore exact numbers cannot be provided per All of Us data dissemination policy.

Activity data were collected from the following laboratory concept names: “Glucose-6-phosphate dehydrogenase (G6PD); quantitative”, “Glucose-6-Phosphate dehydrogenase [Entitic Catalytic Activity] in Blood”, “Glucose-6-Phosphate dehydrogenase [Enzymatic activity/mass] in Red Blood Cells”, “Glucose-6-Phosphate dehydrogenase [Enzymatic activity/volume] in Red Blood Cells”, “Glucose-6-Phosphate dehydrogenase [Enzymatic activity/volume] in Serum”, “Glucose-6-Phosphate dehydrogenase [Presence] in DBS”, “Glucose-6-Phosphate dehydrogenase [Presence] in Red Blood Cells”, and “Glucose-6-Phosphate dehydrogenase [Presence] in Serum”. Values were normalized to the average of the observational medical outcomes partnership (OMOP) standardized lower and upper limits of the normal range provided for each laboratory measurement and multiplied by 100 to give a percent activity of normal. For values that did not have OMOP normal ranges, but were categorized as “normal”, a value of 100% was assigned. For values without ranges or categorization, the value was normalized to the median of all values in that laboratory concept name group. For participants with multiple lab values, we used the lowest value.

### Statistics

P-values for figures were computed in R using t-test or one-way ANOVA with Tukey’s honestly significant difference (HSD) as noted in figure legends. P-values for Table 2 were calculated using Fisher’s exact test in R on 2×2 contingency tables based on allele counts of the variants in alleles from individuals with versus without a diagnosis of G6PD deficiency or G6PD deficiency anemia.

**Table 2:** Missense variant enrichment in individuals with G6PD deficiency in All of Us.

| variant SNP | p-value |
| --- | --- |
| c.202G>A | $< 2.2 \times 10^{-16}$ |
| c.376A>G | $9.1 \times 10^{-13}$ |
| c.563C>T | $1.1 \times 10^{-5}$ |
| c.968T>C | 0.08 |

Enrichment p-values by two sided Fisher exact tests for missense variant SNPs found in 29 individuals with the code "Deficiency of glucose-6-phosphate dehydrogenase" or "Glucose-6-phosphate dehydrogenase deficiency anemia" in All of Us compared to individuals without either code.

## RESULTS

### An updated catalog of G6PD variants

Through database analysis and literature searches, we identified 1,341 G6PD variants that have been reported in humans (Fig. 1A, Supplemental Table 1). 296 of these variants have been described in publications, most in individuals with G6PD deficiency (75%, 221/296).

**Figure 1:**
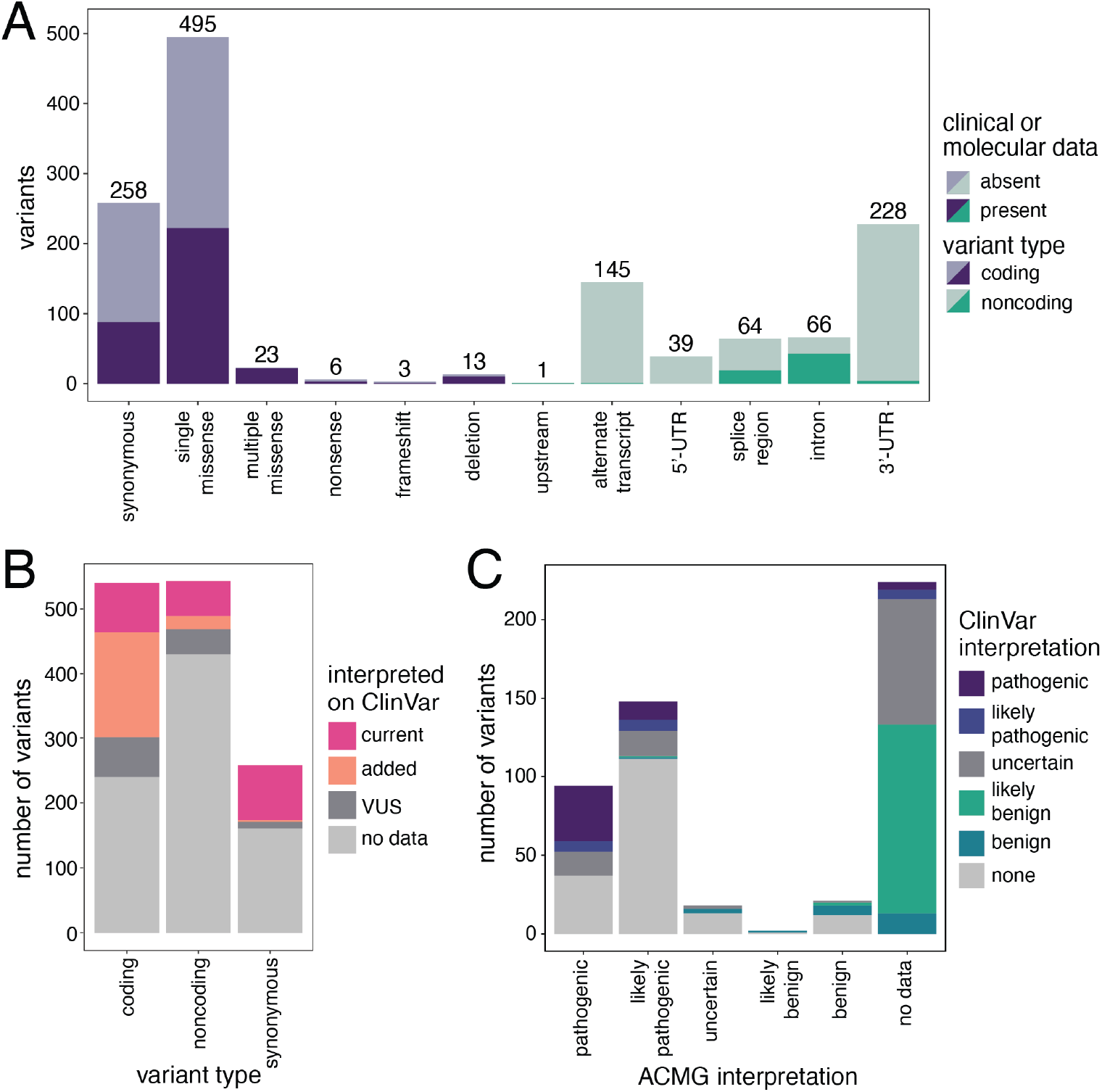
G6PD variant interpretation. (A) A total of 1,341 G6PD variants found from variant database and PubMed queries. Clinical or molecular data indicates the variant is interpreted on ClinVar (as benign, likely benign, likely pathogenic, or pathogenic), or has activity or stability data available from published studies. (B) 214 variants with interpretations currently available on ClinVar, and an additional 186 interpreted by applying ACMG guidelines to published reports. VUS includes variants interpreted as uncertain or with conflicting interpretations, including five previously interpreted on ClinVar. (C) Concordance between variant interpretations on ClinVar and by our application of ACMG guidelines for 509 total G6PD variants.

For 414 variants, additional detailed information on molecular function or clinical interpretation was available (Fig. 1A). However, only 27% (113/414) have been characterized in multiple reports.

### Interpreting G6PD variants using ACMG guidelines

By applying the ACMG standards and guidelines for the interpretation of sequence variants^22,23^ to our collected information, we classified 268 variants, 186 of which were previously unclassified (166) or listed as uncertain significance or other on ClinVar (20) (Fig. 1B, Supplemental Table 4). We also provided interpretations of 12 variants with conflicting interpretations on ClinVar, lending additional support for classification.

We observed a general consensus for 75 variants interpreted on ClinVar with sufficient information for us to also interpret them using ACMG guidelines (Fig. 1C). The 61 variants ClinVar interpreted as pathogenic or likely pathogenic we also interpreted in one of those groups. There were far fewer published reports in support of benign interpretation, but our interpretation agreed with ClinVar for 9/14 benign and likely benign variants. For three variants interpreted on ClinVar as benign, we found conflicting reports of hemizygotes with and without deficiency and wide variation in activity (Supplemental Tables 3 and 4); these conflicts led us to interpret them as variants of uncertain significance. For the other two conflicting variants, one was deposited as likely benign but without evidence to assess independently (variation ID 1546331), while we found a report of decreased activity and G6PD deficiency^33^; the other conflicting variant was deposited to ClinVar by OMIM as benign for its “nearly normal properties” (variation ID 10362), but investigating the referenced reports shows that activity was decreased to 20% of normal.^34,35^ Applying ACMG guidelines to the information on these two variants suggested they are likely pathogenic.

By combining the 219 variants already interpreted on ClinVar with our additional interpretation of 186 variants and five benign variants called into question, 400 G6PD variants now have proposed interpretations other than uncertain or conflicting (Fig. 1B). Most are nonsynonymous coding variants (60%, 239/400), and most of those are single missense variants (85%, 202/239).

### Evaluating G6PD activity in the context of variant classifications

Since G6PD deficiency results from insufficient G6PD activity, we used our collected data on G6PD variant activity to assess different strategies that infer or categorize G6PD variant function by activity. Most G6PD variants associated with deficiency alter the amino acid sequence, and few regulatory variants have been characterized.^1^ We confirmed that variants not affecting the coding sequence had significantly higher activity and were rarely associated with anemia (Fig. 2A). Variants altering the coding sequence led to a range of activity and clinical presentations, but all nonsense variants and deletions led to less than 10% normal activity and CNSHA (Fig. 2B).

**Figure 2:**
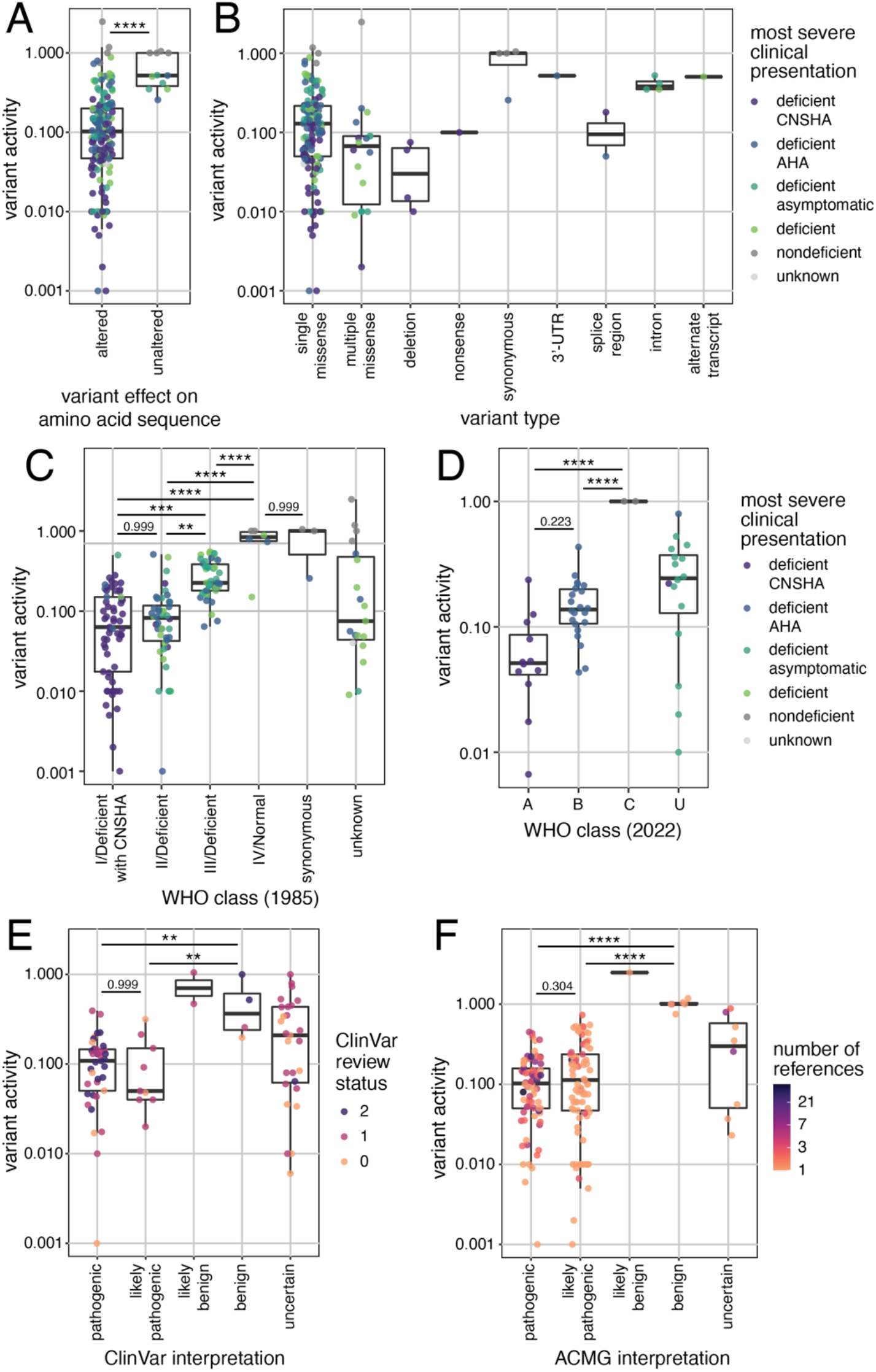
Classification systems for G6PD variants. (A-B) Average G6PD activity by type of genetic alteration in the variant, with most severe reported clinical presentation associated with each variant (“deficiency” indicates further information on anemia was not collected). (C) Average G6PD variant activity by 1985 and (D) 2022 WHO classification, (E) ClinVar interpretation, and (F) by our application of ACMG guidelines. In ClinVar review status, 0 indicates least supporting evidence, and increasing numbers indicate increasing support. All average activities are from red blood cell extracts and weighted by number of hemizygotes per study. P-value by one-way ANOVA with Tukey’s HSD, except A by t-test, ** p<0.01, *** p<0.001, **** p<0.0001.

Using the 1985 WHO classification scheme, we found that G6PD variant activity increases as class levels increase (Fig. 2C), but our data corroborate known issues with the 1985 WHO classification method.^9,10^ Several Class I variants associated with CNSHA have over 10% activity, and for some reported as Class I we were unable to find clinical reports to support the diagnosis of CNSHA. Classes II and III both contained variants that led to AHA, rendering them redundant. Applying the 2022 WHO classification method, which requires variants to have measured activity in at least three unrelated hemizygotes^10^, we observed clearer separation between variants leading to CNSHA and AHA (Fig. 2D). However, many variants that have not been reported to lead to anemia also have activity below 60% of normal and thus were classified as uncertain.

Although the interpretations provided by ClinVar do not require reporting of variant activity, pathogenic and likely pathogenic variants have significantly lower activity than benign variants (Fig. 2E). Variant activity is considered when applying the ACMG guidelines, so pathogenic and likely pathogenic variants have significantly lower activity than benign variants (Fig. 2F).

### G6PD structure impacts variant function and clinical presentation

G6PD contains several domains required for its function, including NADP binding sites, a substrate binding site, and dimerization domain.^26^ The dimerization domain is especially important since G6PD is only active as a dimer and tetramer.^1^ Variants that disrupt the dimerization domain are often predicted to be detrimental, which is evident in our collected data by a cluster of variants in the dimer interface with low activity and stability associated with CNSHA (Fig. 3A-C).

**Figure 3:**
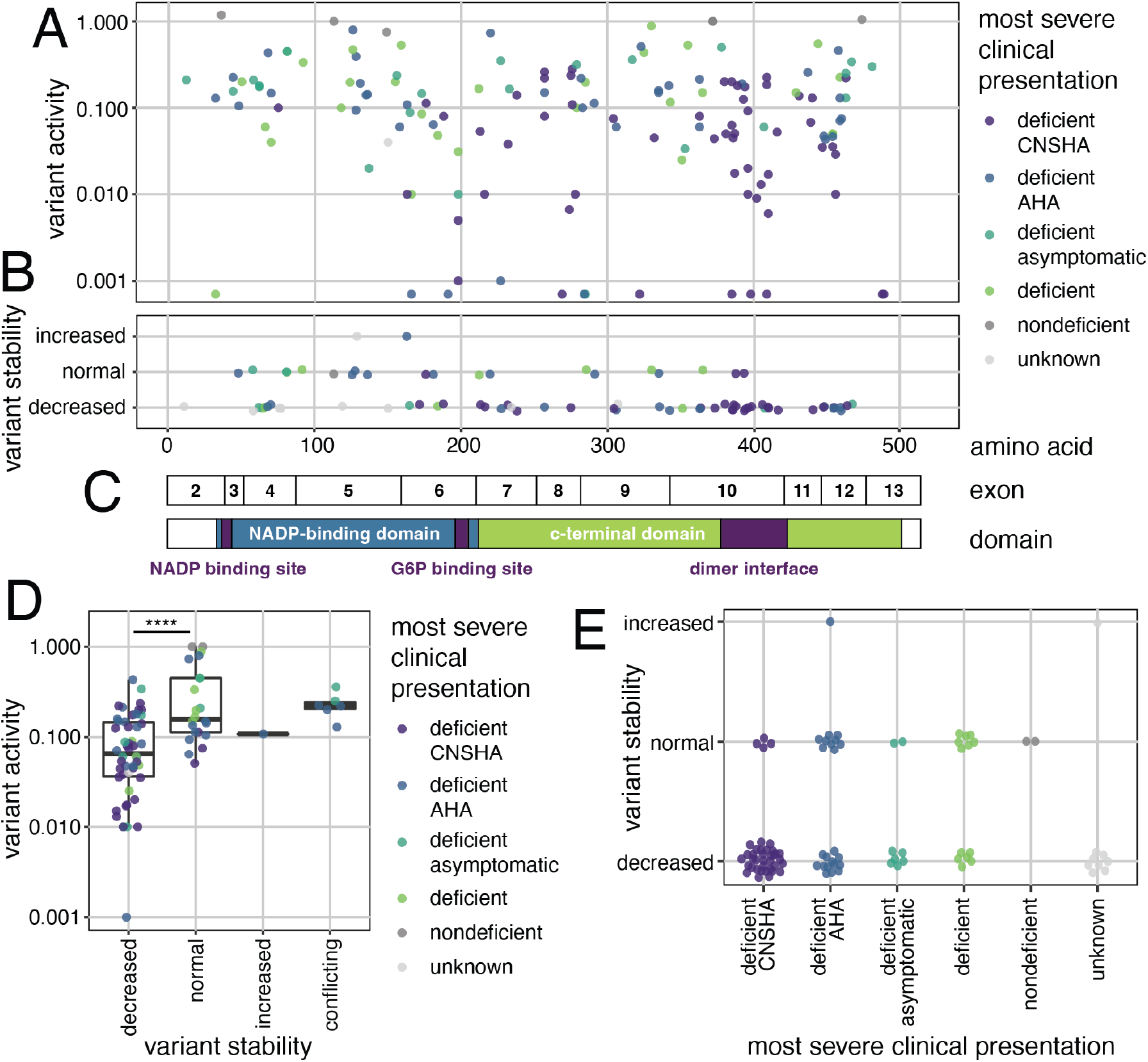
G6PD structure and variant effects. (A) G6PD variant activity and (B) stability across (C) the structural domains, with most severe clinical presentation associated with each variant (“deficiency” indicates further information on anemia was not collected). Only single missense and synonymous variants shown. (D) Variant stability compared to activity and (E) phenotypic severity. All average activities are from red blood cell extracts and weighted by number of hemizygotes per study. P-value by one-way ANOVA with Tukey’s HSD, **** p<0.0001.

Loss of stability is a major mechanism of G6PD deficiency^1^, and we observed that most variants associated with CNSHA were unstable (89%, 34/38) (Fig. 3D-E). Some variants had decreased activity but normal stability, suggesting a decrease in specific activity (Fig. 3D).

### Variation across reports contributes to challenges in variant interpretation

A challenge in interpreting G6PD variant effects is the variability between individuals sharing the same genotype, even when only considering hemizygotes with deficiency.^10^ This was part of the rationale for requiring activity to be measured in three unrelated individuals for the 2022 WHO variant classification, and the variability is clear when observing reported activity for variants with five or more studies (Fig. 4A). Some variants associated with anemia have reports of activity ranging from less than 2% to over 60% of normal (G6PD Chatham, Canton, and Kaiping). Separating variants by clinical presentation still reveals high variability across variants and studies (Fig. 4B), and the activity of variants reported with AHA is not significantly different from ones reported without symptoms.

**Figure 4:**
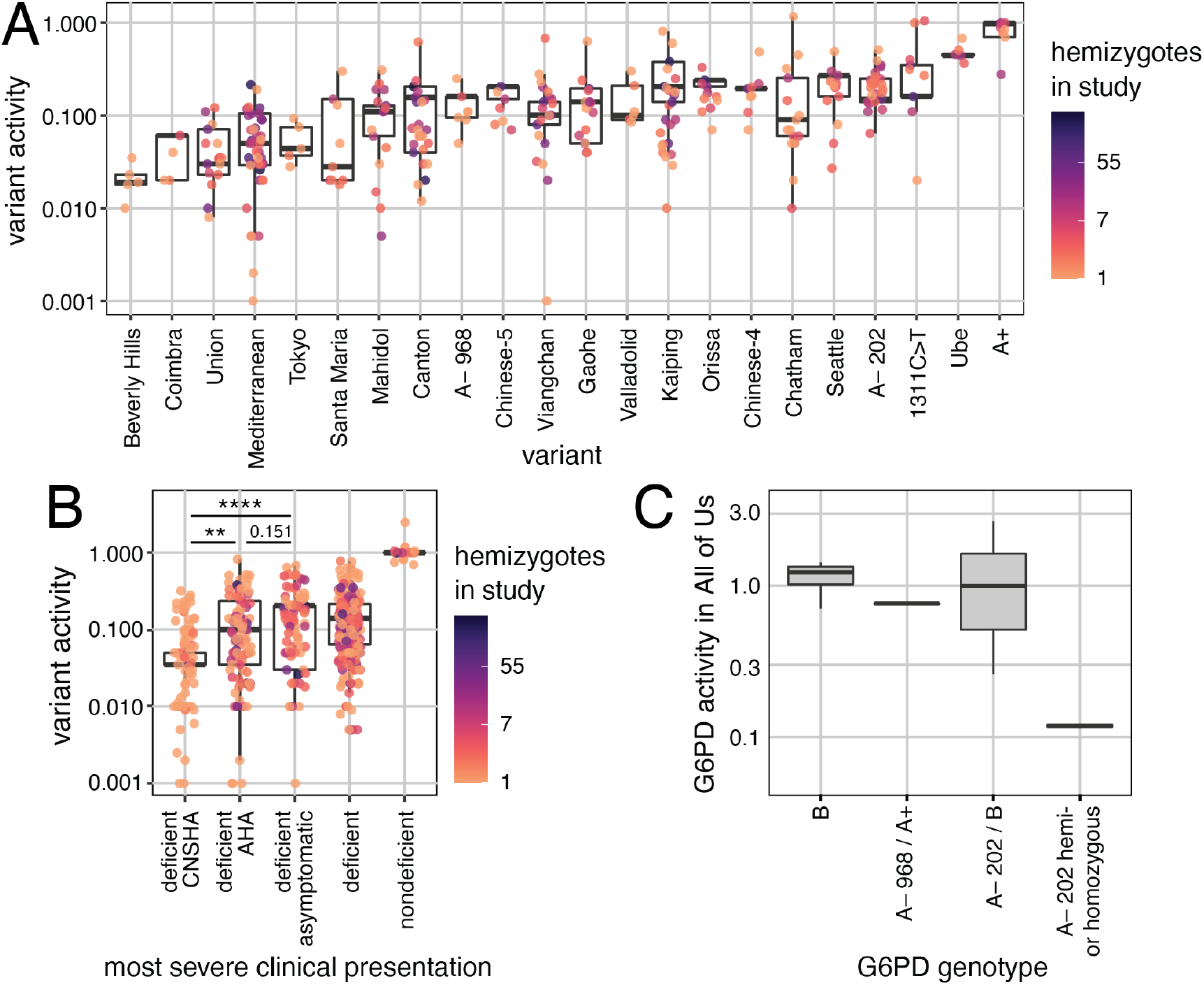
Variation in activity and clinical presentation for G6PD variants across studies. (A) Average G6PD activity (normalized to B variant activity) per study for all variants with at least 5 independent reports. (B) Average G6PD variant activity by most severe clinical presentation in each study. Nondeficient group is significantly different from each other group (p<0.0001). All average activities are from red blood cell extracts of hemizygotes, and box plots weighted by number of hemizygotes per study. P-value by one-way ANOVA with Tukey’s HSD, ** p<0.01, **** p<0.0001. (C) G6PD activity measured in blood or serum for fewer than 20 individuals with the code “Deficiency of glucose-6-phosphate dehydrogenase” or “Glucose-6-phosphate dehydrogenase deficiency anemia” in All of Us. Some individuals with B variant have noncoding variation. Box sizes represent variation between individuals and thus do not reflect the absolute number of individuals.

### Diverse biobank data provide additional support for genotype-phenotype relationships

The majority of G6PD variants with publications supporting their link to deficiency have been reported in fewer than three individuals (70%, 167/240, Supplemental Table 3). The All of Us Research Project has collected clinical codes and whole-genome sequencing from thousands of diverse individuals, enabling us to connect variants with clinical presentations. Whole genome sequencing data included 118 novel variants not identified in any of our prior database or literature searches, including 18 missense and one frameshift variant (Table S1). 29 individuals were reported as diagnosed with G6PD-deficiency, with or without anemia, and carry four G6PD missense variants in different combinations (Table 1 and 2). This includes rarer alleles such as A-968 (c.[376A>G;968T>C]), which has previously only been reported with anemia in three individuals, one homozygote and two heterozygotes.^36,37^ Three missense variants in G6PD (c.202G>A, c.376A>G, and c.563C>T) were significantly enriched in All of Us alleles from participants with a diagnosis of G6PD deficiency (p-values < 0.0001) (Table 2). Occurrence of common A-202 variant c.[202G>A;376A>G] in both G6PD-deficient individuals who are asymptomatic and who have AHA is in agreement with other reports (Supplemental Table 3).

G6PD activity data were available for a subset of the 29 individuals with G6PD deficiency, and agreed with previous reports. Individuals hemizygous or homozygous for the common A-202 variant had decreased activity (average 12% of normal), while heterozygotes had near-normal activity (average 131%) but ranged widely (Fig. 4C). The heterozygous combination of A-968 and A+ variants (c.[376A>G;968T>C];[376A>G]) led to slightly decreased activity (average 76% of normal), similar to activity previously reported in A-968 heterozygotes with deficiency (44-66%).^38^

There were 21 variants present in individuals with a G6PD-deficient anemia diagnosis without any previously known or suspected missense variants causative of G6PD deficiency. These included one synonymous, 15 intronic, and five downstream variants (two in the 3’-UTR). One of the 3’-UTR variants (c.*357G>A) and the synonymous variant (c.1311C>T) have been previously reported with deficiency and anemia in the absence of missense variation, but also in many cases of nondeficiency (Supplementary Table 3). However, despite a diagnosis of G6PD deficiency, individuals without missense variants for whom activity measurements were available did not have decreased G6PD activity (average 115% of normal) (Fig. 4C).

## DISCUSSION

By conducting a systematic review of literature and databases, we compiled a catalog of all G6PD variants reported in humans. Applying the ACMG guidelines to information in published reports enabled us to classify 276 variants, 182 of which were previously unreported or of uncertain significance on ClinVar, bringing the total number of G6PD variants with functional interpretations to 400. We are continuing to maintain an updated catalog of G6PD variants and their effects, available at https://github.com/reneegeck/G6PDcat, so that data on variant function and clinical presentation are available for research and interpretation.

For the variants with data on activity and stability, our collected findings generally support the WHO classification systems that group variants by activity and phenotypic severity^9,10^, but we note that many variants with low activity have not been reported to lead to anemia and thus do not fit the WHO classification thresholds. The data also highlight that while lower variant activity is associated with more severe clinical manifestations, there is a range of activities in each variant class such that one cannot always be accurately predicted from the other. To further confound this, reports of activity and clinical presentation for one variant can vary greatly between individuals and over time.^4^ The 2022 WHO classification system takes inter-individual variation into account, requiring that classification is based on clinical reports of the presence or absence of anemia and the activity from at least three unrelated hemizygotes.^10^ Only 53 variants met these criteria, so we encourage clinicians to continue reporting patients’ variant activity and if they present with anemia. Currently, the variability of activity for many variants emphasizes that point-of-care activity tests remain critical to diagnose deficiency and inform drug prescription and dosage for each individual. Activity tests are also necessary for heterozygotes since their activity varies based on X-inactivation, but knowing that a variant can lead to deficiency in hemizygotes is valuable to support testing in heterozygotes.

Despite efforts to interpret the effects of G6PD variants, the function of most identified variants remains unknown. There is a particular dearth of molecular evidence on variants not associated with anemia or other symptoms. Many of these have been reported through population sequencing, but since individuals with G6PD deficiency can be healthy unless they encounter a trigger, presence of a variant in a population database does not preclude decreased activity. Diverse biobank data, such as that provided by All of Us, enabled us to find 118 novel G6PD variants, 18 of which were rare missense variants. This biobank data will increase our ability to interpret and classify variants, especially when data are provided on G6PD deficiency status or G6PD activity. While most data on G6PD activity and genotype-phenotype correlations have been collected through studies focused on G6PD deficiency, our analysis of All of Us highlights the role that large, diverse biobank data can play in the future of variant interpretation. Recent successes in functionalizing variants with multiplexed assays of variant effect are also being applied to coding variation in G6PD, which will be particularly useful for extremely rare variants with little to no clinical data.^29,39^

While most noncoding variants in G6PD are presumed to be benign or of little effect, this merits further investigation, particularly for variants observed in multiple individuals with G6PD deficiency. Noncoding variants found in individuals with G6PD deficiency but without missense variation, as is the case for some individuals in All of Us, could be a starting point for functional assessment of noncoding variation. However, discordance between diagnostic codes of G6PD deficiency and activity measurements presents a challenge for interpretation. Activity measurements taken during a hemolytic crisis or following a transfusion can be elevated due to high percentages of reticulocytes or donor cells, and lack of that information can confound analysis of biobank data.^4^ Reporting multiple measurements over time compared to nondeficient controls aids in interpretation of G6PD activity tests.^1,4^

Our efforts to classify G6PD variants using published data benefited from the abundant literature, since G6PD deficiency was first described in 1956.^28^ Other genes with considerable variation and many reports with clinical and functional data could be similarly reviewed and analyzed to provide variant interpretations. Determining the gaps between numbers of identified variants and number of characterized variants could highlight genes that would benefit from multiplexed functional studies to provide information on many variants of unknown or uncertain function.

By making use of published studies of G6PD variants, especially those reporting clinical effects, our interpretations can be used to inform clinical decision making. With our addition of 182 interpretations, the number of G6PD variants with data available to aid interpretation has increased to 400 variants. Although there is variability of effects between individuals, adding to the number of variants with proposed interpretations increases the information available to clinicians as they make decisions on how to treat and advise patients with G6PD deficiency. Coupled with robust drug prescribing recommendations^4^, these variant interpretations enable the application of genetic medicine for individuals with G6PD variants.

## Supplemental data

Supplemental data includes five tables.

## Acknowledgements

We thank the following individuals for advice and guidance: S. Fayer, D. Fowler, D. Nickerson, A. Rettie, C. Sibley, A. Stergachis, and members of the Dunham lab. This research was funded in part through the NIH/NCI Cancer Center Support Grant P30 CA015704 and by NIH/NIGMS under award number R01 GM132162. RCG was supported by NIH/NIGMS under award F32 GM143852 and NIH/NHGRI under award T32 HG00035. NRP was supported by NIH/NIGMS under award T32 GM008425.

The All of Us Research Program is supported by the National Institutes of Health, Office of the Director: Regional Medical Centers: 1 OT2 OD026549; 1 OT2 OD026554; 1 OT2 OD026557; 1 OT2 OD026556; 1 OT2 OD026550; 1 OT2 OD 026552; 1 OT2 OD026553; 1 OT2 OD026548; 1 OT2 OD026551; 1 OT2 OD026555; IAA #: AOD 16037; Federally Qualified Health Centers: HHSN 263201600085U; Data and Research Center: 5 U2C OD023196; Biobank: 1 U24 OD023121; The Participant Center: U24 OD023176; Participant Technology Systems Center: 1 U24 OD023163; Communications and Engagement: 3 OT2 OD023205; 3 OT2 OD023206; and Community Partners: 1 OT2 OD025277; 3 OT2 OD025315; 1 OT2 OD025337; 1 OT2 OD025276. In addition, the All of Us Research Program would not be possible without the partnership of its participants.

## Data and code availability

The most current versions of variant lists are available at https://github.com/reneegeck/G6PDcat. Plots, weighted averages and medians, and significance statistics for figures were produced using R code which is also available on GitHub. Code used for All of Us analyses is saved in the All of Us workbook and will be gladly shared with approved users upon request.

## REFERENCES

1. Luzzatto, L., Ally, M., and Notaro, R. (2020). Glucose-6-phosphate dehydrogenase deficiency. Blood 136, 1225–1240.

2. McDonagh, E.M., Thorn, C.F., Bautista, J.M., Youngster, I., Altman, R.B., and Klein, T.E. (2012). PharmGKB summary: very important pharmacogene information for G6PD. Pharmacogenet. Genomics 22, 219–228.

3. Belfield, K.D., and Tichy, E.M. (2018). Review and drug therapy implications of glucose-6-phosphate dehydrogenase deficiency. Am. J. Health-Syst. Pharm. AJHP Off. J. Am. Soc. Health-Syst. Pharm. 75, 97–104.

4. Gammal, R.S., Pirmohamed, M., Somogyi, A.A., Morris, S.A., Formea, C.M., Elchynski, A.L., Oshikoya, K.A., McLeod, H.L., Haidar, C.E., Whirl-Carrillo, M., et al. (2022). Expanded Clinical Pharmacogenetics Implementation Consortium (CPIC) Guideline for Medication Use in the Context of G6PD Genotype. Clin. Pharmacol. Ther. n/a,.

5. DelFavero, J.J., Jnah, A.J., and Newberry, D. (2020). Glucose-6-Phosphate Dehydrogenase Deficiency and the Benefits of Early Screening. Neonatal Netw. NN 39, 270–282.

6. Manjurano, A., Sepulveda, N., Nadjm, B., Mtove, G., Wangai, H., Maxwell, C., Olomi, R., Reyburn, H., Riley, E.M., Drakeley, C.J., et al. (2015). African Glucose-6-Phosphate Dehydrogenase Alleles Associated with Protection from Severe Malaria in Heterozygous Females in Tanzania. PLOS Genet. 11, e1004960.

7. Yoshida, A., Beutler, E., and Motulsky, A.G. (1971). Human glucose-6-phosphate dehydrogenase variants. Bull. World Health Organ. 45, 243–253.

8. WHO Working Group (1989). Glucose-6-phosphate dehydrogenase deficiency. Bull. World Health Organ. 67, 601–611.

9. Malaria Policy Advisory Committee (2019). Updating the WHO G6PD classification of variants and the International Classification of Diseases, 11th Revision (ICD-11) (World Health Organization).

10. Malaria Policy Advisory Committee (2022). Technical consultation to review the classification of glucose-6-phosphate dehydrogenase (G6PD) (World Health Organization).

11. Relling, M.V., McDonagh, E.M., Chang, T., Caudle, K.E., McLeod, H.L., Haidar, C.E., Klein, T., and Luzzatto, L. (2014). Clinical Pharmacogenetics Implementation Consortium (CPIC) Guidelines for Rasburicase Therapy in the Context of G6PD Deficiency Genotype. Clin. Pharmacol. Ther. 96, 169–174.

12. Landrum, M.J., Lee, J.M., Benson, M., Brown, G.R., Chao, C., Chitipiralla, S., Gu, B., Hart, J., Hoffman, D., Jang, W., et al. (2018). ClinVar: improving access to variant interpretations and supporting evidence. Nucleic Acids Res. 46, D1062–D1067.

13. Sherry, S.T., Ward, M.-H., Kholodov, M., Baker, J., Phan, L., Smigielski, E.M., and Sirotkin, K. (2001). dbSNP: the NCBI database of genetic variation. Nucleic Acids Res. 29, 308–311.

14. Karczewski, K.J., Francioli, L.C., Tiao, G., Cummings, B.B., Alföldi, J., Wang, Q., Collins, R.L., Laricchia, K.M., Ganna, A., Birnbaum, D.P., et al. (2020). The mutational constraint spectrum quantified from variation in 141,456 humans. Nature 581, 434–443.

15. Minucci, A., Moradkhani, K., Hwang, M.J., Zuppi, C., Giardina, B., and Capoluongo, E. (2012). Glucose-6-phosphate dehydrogenase (G6PD) mutations database: review of the “old” and update of the new mutations. Blood Cells. Mol. Dis. 48, 154–165.

16. Stenson, P.D., Ball, E.V., Mort, M., Phillips, A.D., Shiel, J.A., Thomas, N.S.T., Abeysinghe, S., Krawczak, M., and Cooper, D.N. (2003). Human Gene Mutation Database (HGMD): 2003 update. Hum. Mutat. 21, 577–581.

17. Rangwala, S.H., Kuznetsov, A., Ananiev, V., Asztalos, A., Borodin, E., Evgeniev, V., Joukov, V., Lotov, V., Pannu, R., Rudnev, D., et al. (2021). Accessing NCBI data using the NCBI Sequence Viewer and Genome Data Viewer (GDV). Genome Res. 31, 159–169.

18. Drousiotou, A., Touma, E.H., Andreou, N., Loiselet, J., Angastiniotis, M., Verrelli, B.C., and Tishkoff, S.A. (2004). Molecular characterization of G6PD deficiency in Cyprus. Blood Cells. Mol. Dis. 33, 25–30.

19. Menounos, P., Zervas, C., Garinis, G., Doukas, C., Kolokithopoulos, D., Tegos, C., and Patrinos, G.P. (2000). Molecular heterogeneity of the glucose-6-phosphate dehydrogenase deficiency in the Hellenic population. Hum. Hered. 50, 237–241.

20. Arunachalam, A.K., Sumithra, S., Maddali, M., Fouzia, N.A., Abraham, A., George, B., and Edison, E.S. (2020). Molecular Characterization of G6PD Deficiency: Report of Three Novel G6PD Variants. Indian J. Hematol. Blood Transfus. Off. J. Indian Soc. Hematol. Blood Transfus. 36, 349–355.

21. Edison, E.S., Melinkeri, S.R., and Chandy, M. (2006). A novel missense mutation in glucose-6-phosphate dehydrogenase gene causing chronic nonspherocytic hemolytic anemia in an Indian family. Ann. Hematol. 85, 879–880.

22. Richards, S., Aziz, N., Bale, S., Bick, D., Das, S., Gastier-Foster, J., Grody, W.W., Hegde, M., Lyon, E., Spector, E., et al. (2015). Standards and guidelines for the interpretation of sequence variants: a joint consensus recommendation of the American College of Medical Genetics and Genomics and the Association for Molecular Pathology. Genet. Med. Off. J. Am. Coll. Med. Genet. 17,.

23. Tavtigian, S.V., Greenblatt, M.S., Harrison, S.M., Nussbaum, R.L., Prabhu, S.A., Boucher, K.M., Biesecker, L.G., and ClinGen Sequence Variant Interpretation Working Group (ClinGen SVI) (2018). Modeling the ACMG/AMP variant classification guidelines as a Bayesian classification framework. Genet. Med. Off. J. Am. Coll. Med. Genet. 20, 1054–1060.

24. Minucci, A., Concolino, P., Antenucci, M., Santonocito, C., Ameglio, F., Zuppi, C., Giardina, B., and Capoluongo, E. (2007). Description of a novel missense mutation of glucose-6-phosphate dehydrogenase gene associated with asymptomatic high enzyme deficiency. Clin. Biochem. 40, 856–858.

25. Mehta, A., Mason, P.J., and Vulliamy, T.J. (2000). Glucose-6-phosphate dehydrogenase deficiency. Baillieres Best Pract. Res. Clin. Haematol. 13, 21–38.

26. Chiu, Y.-H., Liu, Y.-N., Chen, H.-J., Chang, Y.-C., Kao, S.-M., Liu, M.-Y., Weng, Y.-Y., Hsiao, K.-J., and Liu, T.-T. (2019). Prediction of functional consequences of the five newly discovered G6PD variations in Taiwan. Data Brief 25, 104129.

27. Nkhoma, E.T., Poole, C., Vannappagari, V., Hall, S.A., and Beutler, E. (2009). The global prevalence of glucose-6-phosphate dehydrogenase deficiency: a systematic review and meta-analysis. Blood Cells. Mol. Dis. 42, 267–278.

28. Howes, R.E., Piel, F.B., Patil, A.P., Nyangiri, O.A., Gething, P.W., Dewi, M., Hogg, M.M., Battle, K.E., Padilla, C.D., Baird, J.K., et al. (2012). G6PD deficiency prevalence and estimates of affected populations in malaria endemic countries: a geostatistical model-based map. PLoS Med. 9, e1001339.

29. Fayer, S., Horton, C., Dines, J.N., Rubin, A.F., Richardson, M.E., McGoldrick, K., Hernandez, F., Pesaran, T., Karam, R., Shirts, B.H., et al. (2021). Closing the gap: Systematic integration of multiplexed functional data resolves variants of uncertain significance in BRCA1, TP53, and PTEN. Am. J. Hum. Genet. 108, 2248–2258.

30. Vulliamy, T., Luzzatto, L., Hirono, A., and Beutler, E. (1997). Hematologically important mutations: glucose-6-phosphate dehydrogenase. Blood Cells. Mol. Dis. 23, 302–313.

31. Numanagić, I., Malikić, S., Ford, M., Qin, X., Toji, L., Radovich, M., Skaar, T.C., Pratt, V.M., Berger, B., Scherer, S., et al. (2018). Allelic decomposition and exact genotyping of highly polymorphic and structurally variant genes. Nat. Commun. 9, 828.

32. Hari, A., Zhou, Q., Gonzaludo, N., Harting, J., Scott, S.A., Sahinalp, S.C., and Numanagić, I. (2022). Aldy 4: An efficient genotyper and star-allele caller for pharmacogenomics. BioRxiv.

33. Vaca, G., Arámbula, E., Monsalvo, A., Medina, C., Nuñez, C., Sandoval, L., and López-Guido, B. (2003). Glucose-6-phosphate dehydrogenase (G-6-PD) mutations in Mexico: four new G-6-PD variants. Blood Cells. Mol. Dis. 31, 112–120.

34. Viglietto, G., Montanaro, V., Calabrò, V., Vallone, D., D’Urso, M., Persico, M.G., and Battistuzzi, G. (1990). Common glucose-6-phosphate dehydrogenase (G6PD) variants from the Italian population: biochemical and molecular characterization. Ann. Hum. Genet. 54, 1–15.

35. Calabrò, V., Giacobbe, A., Vallone, D., Montanaro, V., Cascone, A., Filosa, S., and Battistuzzi, G. (1990). Genetic heterogeneity at the glucose-6-phosphate dehydrogenase locus in southern Italy: a study on a population from the Matera district. Hum. Genet. 86, 49–53.

36. Vulliamy, T., Rovira, A., Yusoff, N., Colomer, D., Luzzatto, L., and Vives-Corrons, J.L. (1996). Independent origin of single and double mutations in the human glucose 6-phosphate dehydrogenase gene. Hum. Mutat. 8, 311–318.

37. Benmansour, I., Moradkhani, K., Moumni, I., Wajcman, H., Hafsia, R., Ghanem, A., Abbès, S., and Préhu, C. (2013). Two new class III G6PD variants [G6PD Tunis (c.920A>C: p.307Gln>Pro) and G6PD Nefza (c.968T>C: p.323 Leu>Pro)] and overview of the spectrum of mutations in Tunisia. Blood Cells. Mol. Dis. 50, 110–114.

38. Powers, J.L., Best, D.H., and Grenache, D.G. (2018). Genotype–Phenotype Correlations of Glucose-6-Phosphate–Deficient Variants Throughout an Activity Distribution. J. Appl. Lab. Med. 2, 841–850.

39. Geck, R.C., Boyle, G., Amorosi, C.J., Fowler, D.M., and Dunham, M.J. (2022). Measuring Pharmacogene Variant Function at Scale Using Multiplexed Assays. Annu. Rev. Pharmacol. Toxicol. 62, 531–550.

